# From Hands to Feet: Experience-Driven Plasticity in Secondary Somatosensory Cortex in People Born without Hands

**DOI:** 10.64898/2025.12.09.693253

**Authors:** Zhiqing Deng, Florencia Martinez-Addiego, Yuqi Liu, Ella Striem-Amit

## Abstract

In congenital handlessness, where individuals rely on their feet for compensatory manual functions, reorganization in the primary somatosensory cortex (SI) appears limited to local shifts by adjacent body parts. It remains unclear how congenital handlessness and lifelong compensatory foot use shape the sensorimotor system beyond SI, and whether compensatory abilities can drive broader functional reorganization. Here, we used task-based and resting-state functional magnetic resonance imaging (fMRI) to investigate how compensatory foot use changes body representations in the secondary somatosensory cortex (SII) in people born without hands (individuals with upper-limb dysplasia, IDs). We found that IDs showed foot selectivity in the left hand-selective SII, with stronger foot activity than typically-developed (TD) control participants during action execution. Notably, this reorganization in the left SII hand area cannot simply be explained by this area’s intrinsic organization - the typical secondary preference of this area in the TDs when hand responses are excluded, as this area in TDs shows shoulder - but not foot - selectivity when the hand activity is excluded. This indicates a functional reorganization dependent on the IDs’ sensorimotor experience. Resting-state results further revealed altered functional connectivity between SII and primary sensorimotor regions in IDs. Our findings suggest that SII exhibits flexible, experience-dependent plasticity, revealing a hierarchical principle of cortical plasticity, whereby reorganization in the SI is constrained by somatotopic principles while SII reflects functionally or experience-dependent plasticity.

## Introduction

Somatotopy - the ordered mapping of body parts such that adjacent body parts are represented in adjacent cortical locations - is a fundamental organizing principle in primary motor (M1) and somatosensory cortices (SI) [1, 2]. In both cortices, the hand-responsive area takes a large share due to their behavioral relevance in typically developed individuals (TDs), who use their hands as their main body part for everyday actions. Unlike them, people born without both hands (individuals with upper-limb dysplasia, IDs) rely on their feet for manual actions, providing a unique opportunity to study how congenital hand absence reshapes brain organization. Although the missing hand territory in SI/M1 is activated by many body parts in people born without one hand or born with upper extremity dysmelia [3, 4], it is preferentially taken over by adjacent body-part representations in IDs, such as the shoulder, indicating that reorganization in the primary sensorimotor cortex is constrained by somatotopic principles [5]. Similar somatotopic constraints have been observed in adult humans and monkeys with digit or arm amputation, where, to the degree that reorganization occurs in SI/M1 (SM1) ([6, 7]), it is limited to the immediate topographic neighboring body-part representations that invade deprived territory [6-9]).

In contrast to the primary sensorimotor cortex, higher-order association cortices such as the parietal cortex receive information from SI/M1, integrate somatosensory and visual information, and link perception and action [10, 11]. Unlike primary areas, these regions appear to be organized more by function than by strict somatotopic sensory input: Recent evidence indicates that higher-level sensorimotor areas encode more abstract action representations, which extend across acting body parts (effectors; [12-20]). These areas exhibit the same preference for the performed actions, even in individuals born without hands, who perform these actions with their feet [12, 13]. Thus, while SI/M1 reorganization appears to be constrained by somatotopy, the higher-order association cortex may support more flexible, functionally driven plasticity. Consistent with this idea, we previously found that in IDs, the anterior intraparietal sulcus (aIPS) and inferior parietal lobule (IPL) showed foot selectivity, suggesting experience-driven or functionally driven plasticity in higher-order association cortices [5].

How does this reorganization occur? Where in the sensorimotor hierarchy is there re-wiring that changes from a topographical to a functional code? As an intermediate processing hub, interacting with both SI and higher-order parietal areas [21-23], the secondary somatosensory cortex (SII) is known to respond to somatosensory stimuli [24], characterized by larger, bilateral, and overlapping receptive fields [24-29]. While traditionally considered to show a similar pattern to SI in higher primates [30-32], SII integrates somatosensory information [33] and also transmits information to the parietal cortex [22], suggesting a broader integrative role than SI. Therefore, we ask whether it may serve as a transition between functional codes: if its organization is limited by somatotopic principles, or if it shows flexible and experience-driven or functionally dependent plasticity in people born without hands.

Here, we used task-based and resting-state functional magnetic resonance imaging (fMRI) to investigate how congenital handlessness and compensatory foot use change body representation in SII and its functional connectivity in people born without hands. In TDs, we defined hand-selective regions across the brain by contrasting right-hand activity and activity for other body parts. Next, we examined body-part selectivity in “missing hand” territory (hand-selective regions in TDs) in people born without hands to test whether congenital hand absence induces reorganization in SII constrained by intrinsic organization or reflects experience-dependent functional plasticity. We also tested whether such reorganization generalized to different motor execution tasks. Last, we examined functional connectivity between SI/M1 and SII to assess whether congenital handlessness shapes functional coupling within the sensorimotor system. Together, these analyses test the functional plasticity of the human somatosensory system.

## Results

We investigated how congenital hand absence and compensatory experiences shape brain organization beyond the primary somatosensory cortex by combining task-based and resting-state fMRI in people born without hands (IDs) and typically developed individuals (TDs). Participants completed two fMRI experiments: Experiment 1: Motor Mapping Experiment, where they performed simple flexing movements of different body parts (including hands for TDs, feet, shoulders, abdomen, and mouth) in different blocks, and Experiment 2: Action Experiment, where they performed object-directed actions using their hands (only TDs) or feet (both TDs and IDs). Four kinds of object-directed actions were included: grasping, simple tool-use, drawing, and writing. In addition, resting-state fMRI data were also collected to reveal the functional connectivity among somatosensory cortices.

First, hand-selective brain regions were defined in TDs by comparing right hand and other body part activities in the motor mapping experiment (Experiment 1; *p*_*corr*_ < 0.05, see **Fig. 1a)**. Within these areas, we examined the body part selectivity in IDs using the contrast between each body part and the others (see **Fig. 1b**). Consistent with previous work [5], the missing hand territory in SI/M1 (SM1) was selectively taken over by nearby body parts (shoulder, abdomen, mouth, see **Fig. 1b**) in IDs. These results further support the idea that reorganization in the primary somatosensory cortex is constrained by somatotopic principles.

**Figure 1.**
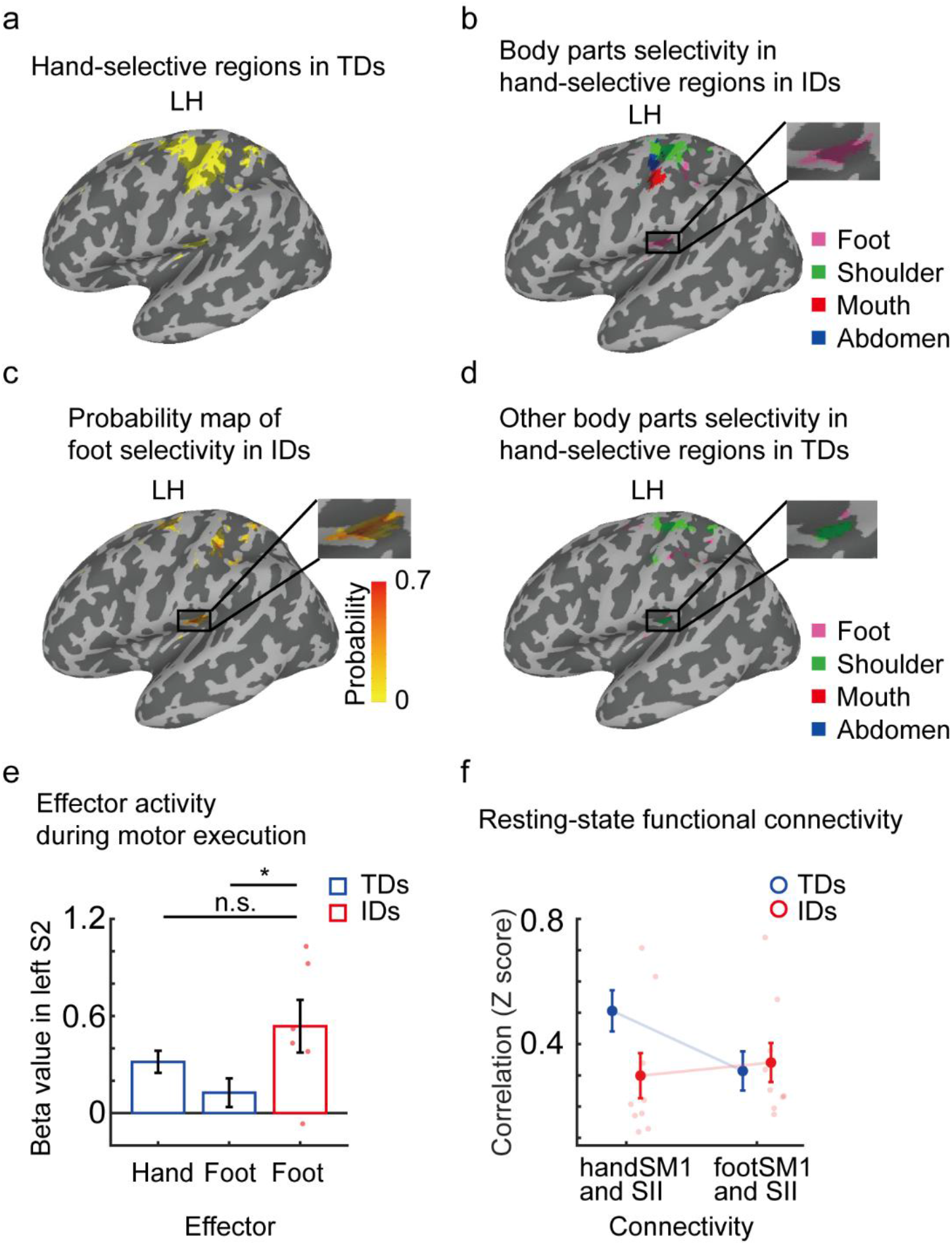
**(a)** Hand-selective regions were defined in typically developed individuals (TDs) by contrasting between the right hand and other body parts (right foot, right shoulder, stomach, and mouth) activity. **(b)** Body part selectivity analysis in the hand-selective regions was applied in people born without hands (individuals with upper-limb dysplasia, IDs) by contrasting each body part activity with the other body parts. **(c)** To test the consistency of foot selectivity across IDs, an overlap map was calculated for foot selectivity in hand-selective regions across individual maps of all IDs. Sixty-six percent of them (corresponding to six of the nine participants) showed significant foot selectivity in left hand-selective SII. **(d)** A body-part selectivity analysis excluding the hand, same as that in IDs, was also applied in TDs, to test if the SII hand area shows a secondary preference for the foot. This was calculated by contrasting each body part activity with the other body parts, except the right hand. **(e)** Comparison between hand or foot activity in TDs and foot activity in IDs for goal-directed actions (Experiment 2). This was statistically assessed using bootstrapping methods due to small sample size of IDs group. Significance was determined by nonparametric bootstrapping (10,000 iterations). * *p* < 0.05. **(f)** Seed-based functional connectivity between hand-selective SII and hand or foot-selective SM1 (handSM1 or footSM1) using resting-state fMRI data shows a significant change in connectivity between the groups (Group × Connectivity interaction).

More strikingly, beyond SI/M1, the left hemisphere hand-selective secondary somatosensory cortex (SII) showed a preference for foot activity in IDs (*pcorr* < 0.05, **Fig. 1b**). Importantly, this pattern was consistent across individuals: most IDs (6 out of 9) showed foot selectivity in left SII (**Fig. 1c**). To verify our results do not stem from interindividual variability in the location of the SII missing-hand area, we inspected the body-part preference in the entire SII at the single-subject level. For this, we calculated winner-take-all maps in the left SII (Area OP1, parietal operculum 1, defined using a cytoarchitectonic atlas [34]). Consistent with body selectivity in hand-selective SII in TDs (**Fig. 1a**), and with the known somatotopic organization of this region [26, 35], a large part of SII responded selectively to the hands in TDs, with mouth-selective territories appearing along the superior and lateral margin of the SII region (**Fig. S1a**). Smaller preferential responses to the shoulder and feet were found in the posterior and inferior portion of SII in TDs (**Fig. S1a**). However, in the IDs, the majority of SII - overlapping with the hand area in TDs - responded selectively to the foot (**Fig. S1b**). Importantly, individual winner-take-all maps in IDs demonstrated that foot preference predominantly took over the SII central region for most participants (**Fig. S1c**; 8 out of 9 participants).

Theoretically, one might argue that this could stem from the intrinsic organization of SII in TDs, in which a foot preference might emerge when hand responses are excluded, if the area’s secondary preference is for the feet (e.g. see discussion in [7]). To test this alternative explanation, we performed the same body parts selectivity analysis in TDs while excluding the hand condition, matching the analysis in IDs. In TDs, hand-selective SII showed a preference for the shoulder, but not for the foot, in this analysis for both SI and SII (see **Fig. 1d**). This indicates that while shoulder remapping in the missing-hand area in SI in IDs may reflect intrinsic organization observed in TDs, the foot selectivity in SII could not simply be explained by the intrinsic organization in TDs and therefore reflect experience-driven reorganization.

We then tested whether this reorganization was consistent during different motor actions by inspecting object-directed actions performed by the hand (only TDs) and foot in another experiment. During motor execution in the Action Experiment (Experiment 2), in left hand-selective SII, response to foot activity in IDs was significantly higher than for foot activity in TDs (mean beta values and standard error: foot activity in IDs: 0.537 ± 0.163, foot activity in TDs: 0.126 ± 0.089, mean difference: -0.410; 95% bootstrap CI [-0.773, -0.055]), 10,000 resamples; Bayes Factor = 4.449). There was no significant difference between foot activity in IDs and hand activity in TDs in this area (mean beta values: foot activity in IDs: 0.537 ± 0.163, hand activity in TDs: 0.317 ± 0.068, mean difference: -0.220, 95% bootstrap CI [-0.566, 0.124], 10,000 resamples; Bayes Factor = 1.515, see **Fig. 1e**).

Our findings indicate that in people born without hands, the missing hand territory in SII is reorganized into foot selectivity, reflecting experience-driven plasticity from the hands to the feet, across simple flexing actions and goal-oriented actions. Hand-selective SII significantly reorganizes for the feet in IDs, and this reorganization is not due to an intrinsic organization in TDs. Instead, it reflects experience-driven reorganization.

For SII to reorganize to a non-adjacent body part while SI maintains nearby topographic preferences implies that experience may alter not only the local representation in SII, but also its connectivity with SI/M1. To test this hypothesis, we calculated the seed-based functional connectivity between left hand-selective SII and hand or foot-selective S1/M1 (handSM1 or footSM1; defined in the TDs group) using resting-state fMRI data. A repeated-measure Bayesian ANOVA results showed strong evidence supporting the main effects of connectivity, as well as, importantly, its interaction with the group factor: a change in connectivity between the seeds across the groups (Bayes Factor _(Model)_ = 13.947; main effect of connectivity and interaction between connectivity and group: both Bayes Factor > 99.851, **Fig. 1f**). In TDs, Bayesian t-tests provided strong evidence that left hand-selective SII was more strongly connected with handSM1 than footSM1 (Bayes Factor = 1605.070, **Fig. 1f**, blue markers). With congenital hand absence (IDs), however, there is no evidence for a difference in between the connectivity of handSM1 and left hand-selective SII and connectivity of footSM1 and left hand-selective SII (Bayes Factor = 0.541, **Fig. 1f**, red markers). Our findings indicate that congenital hand absence alters not only body-part responses within SII, but also shapes its coupling with primary motor and somatosensory cortex.

## Discussion

In this study, we combined task-based and resting-state fMRI data to investigate how congenital hand absence and motor experience shape the organization in the secondary somatosensory cortex. Interestingly, we found IDs showed foot selectivity in a typically hand-selective area of left SII (**Fig. 1b, c**, and **e**). This pattern was reproducible across participants and robust across tasks: the same SII territory showed stronger response to foot activity in IDs in both body-part mapping and independent action execution tasks (**Fig. 1e**). Notably, this reorganization in SII missing-hand territory in IDs cannot simply be explained by the intrinsic organization in TDs, as the secondary selectivity of this region in TDs is for the shoulder, and not the foot (**Fig. 1d**). Instead, this reflects a functional reorganization dependent on the IDs’ sensorimotor experience. Our findings suggest that the secondary somatosensory cortex exhibits experience-dependent plasticity.

The pattern of reorganization in SII diverges sharply from that of the primary somatosensory cortex (SI). In IDs, missing hand territories in SI and M1 are predominantly taken over by neighboring body parts such as the shoulder, abdomen, or mouth (**Fig. 1b**; note that our data is not of sufficient resolution to differentiate between S1 and M1). This finding is consistent with the classical view that primary maps reorganize only across somatotopic boundaries, for both cases of late-onset amputation [9] and congenitally missing hands [5]. In IDs, while the missing-hand territory in SI/M1 was primarily taken over by the shoulder, we also observed some selectivity for responses from the mouth, consistent with findings in amputees showing that the face may invade missing hand territory in SI [9]. Importantly, this mouth takeover was secondary to the robust shoulder takeover in the central missing-hand area, and is consistent with the overall somatotopic reorganization for adjacent body parts observed in SI/M1. Recently, even the ability to reorganize within SI in late-onset amputation was doubted, showing stable cortical representations of both hand and lips in primary sensorimotor regions [7]. This highlights the relative stability and overall constraints on reorganization in SI, both early in life, and especially following amputation in adulthood.

Although SII has also been reported to show somatotopic reorganization [36], it exhibits a more integrative organization than SI: it possesses larger, often bilateral receptive fields and integrates somatosensory input across effectors [24-29]. This architecture likely underlies its capacity for reorganization beyond somatotopic constraints, aligning with the idea that higher cortical areas have more potential for reorganization [37]. The present results suggest that while SI reorganization follows intrinsic topography (see **Fig. 1b, d**; similar to [5]), SII plasticity is guided by functional and experience-driven principles. Note, this result cannot be simply explained by interindividual variability during SII definition: consistent with the main finding (**Fig. 1b-c, Fig. S1a-b**), the missing-hand SII area (hand-selective SII in TDs) was remapped to the foot in the majority of IDs, showing functionally driven plasticity. This idea is consistent with past evidence of functional adaptations in SII following both training and sensory input loss. Tool-use training modifies SII structure and function in monkeys [38], and SII regions can be recruited by alternative effectors after a late-onset SI lesion [37, 39]. Its recruitment during movement preparation is also dependent on one’s prior motor experience [40]. Extending this framework, our findings demonstrate that congenital hand absence drives functionally-driven plasticity in SII.

Our resting-state analyses extend this finding to the network level, showing that congenital hand absence not only changes the local representations in SII, but also shapes functional connectivity within the sensorimotor network. In TDs, the hand-selective region of SII showed stronger connectivity with the hand areas of SI/M1 than with the foot areas, reflecting canonical coupling between corresponding somatotopic territories. In IDs, this distinction was absent: the connectivity between SII and the SM1 hand area weakened as compared to its connectivity to the SM1 foot area (**Fig. 1f**). This reorganization joins altered connectivity observed in congenital one-handers and amputees [41, 42], and highlights that congenital limb absence reshapes not only local responses but also long-range interactions within the sensorimotor network. Such changes are also consistent with broader evidence that SII connectivity is highly experience-dependent and changes during motor learning [43, 44].

Together, these results support a **hierarchical principle of cortical plasticity**. In the primary sensorimotor cortex, reorganization remains constrained by somatotopic principles, preserving stable intrinsic local maps (**Fig. 1b** and **d**) despite altered input. In contrast, higher-order regions such as SII reorganize according to functional demands and experience, integrating the compensatory effector into existing sensorimotor networks. This hierarchical gradient mirrors observations in other sensory systems: in the visual domain, in congenital achromatopsia, V1 maintains highly stable retinotopic maps despite lifelong absence of cone input, whereas higher-order visual regions, such as V3, display marked plasticity that may optimize the use of the available information [45]. Across modalities, it appears that the cortex compensates for partial loss, such as the loss of a body part or part of the perceptual space, by **balancing stability at lower levels with flexibility at higher ones**. Primary sensory areas preserve a relatively stable organization constrained by somatotopic maps to prevent coding disruptions, while higher-order areas flexibly reorganize to optimize behavior under altered input conditions [46]. An alternative conceptual framing of such hierarchical effects, which is not mutually exclusive, is that reorganization is constrained **by their typical functional roles and connectivity** [13]. Higher-level stations in the sensorimotor hierarchy that code for complex action plans and their performance, but may generalize across body parts even in the typical brain, may therefore be more flexible and able to switch between dominant body parts in case of loss. In contrast, earlier stages of the motor hierarchy, like the primary sensorimotor cortex, may be more tied to percepts from specific receptor spaces or specific muscles, which limits their ability to adapt. This is consistent with observations in people born without hands, where actions like reaching, grasping, and tool-use recruit the typical areas in the association motor system [12, 47]. Future studies may also inspect how these areas, specifically cortical premotor, parietal, and non-cortical structures (cerebellum, basal ganglia) shape their somatotopic preferences to allow such compensatory adaptations. However, overall, this hierarchical organization in the motor system helps to stabilize and facilitate goal-directed actions [48-50] and motor generation and learning [51, 52].

Functionally, reallocating deprived SII hand territory to the foot may preserve tactile-motor integration required for planning and executing goal-directed actions. SII is an intermediate stage interacting with both SI and parietal cortex [21-23], contributing to tactile perception and object recognition [53-55] and memory [29, 56-58]. SII also interacts with ventral premotor and prefrontal cortices [59], supporting context-dependent modulation of somatosensory representations during decision making [56, 60]. The observed experience-driven SII plasticity in our study preserves the broader motor hierarchy required for sustaining goal-directed behaviors even when the primary effector is absent.

This functional role is supported by SII’s developmental and evolutionary properties. SII, situated in the subdivision of the parietal operculum (OP1), is a later-evolving and more plastic cortical region [37, 61-63], than the primary somatosensory cortex. Unlike the evolutionarily conserved SI, SII’s circuitry allows dynamic adaptation to environmental demands, providing an extended window for experience-dependent modification and enabling continued functional integration after congenital limb absence. This potential for developmental reorganization likely acts through physiological mechanisms such as Hebbian plasticity - experience-dependent strengthening and weakening of pre-existing synaptic connections [64] - and postnatal axonal remodeling and synapse relocation observed in animal models of deprivation remodeling [65, 66]. This is combined with an evolutionary preferential connectivity pattern, as hand-selective SII may have the capacity to reorganize for the foot glabrous skin [39]. In this light, the altered SII connectivity in congenital hand absence may reflect a developmental analogue of experience-dependent learning, arising from atypical sensory–motor experience across development.

In sum, our findings demonstrate that congenital hand absence induces hierarchically organized plasticity in the sensorimotor system: SI/M1 reorganization is constrained by somatotopic principles, while SII and higher-level regions show experience-driven plasticity. This hierarchical principle preserves the stability of foundational maps while enabling functional adaptation at higher levels, offering a global framework for understanding cortical plasticity in congenital limb absence.

## Materials and Methods

### Participants

Twenty-two typically developed control participants (11 females; mean age: 31.1; age range: 20-65, abbreviated TDs) and nine individuals born with severely shortened or completely absent upper limbs (individuals with upper-limb dysplasia; abbreviated IDs; four females; mean age: 38; age range: 21-62) participated in the experiment. Data from all IDs had been reported in previous neuroimaging and behavioral studies [5, 12, 13, 67-70]. Importantly, all recruited IDs perform everyday actions primarily using their feet. Eight IDs were right-footed, and one ID (ID 6) was left-footed. Since the majority of ID participants are right-footed, controls were required to be right-handed and right-footed (handedness/footedness was determined by self-report, which has been shown to strongly correspond to experimental measures; [71]).

Seventeen TDs and 9 IDs participated in Experiment 1 (Motor mapping experiment). 18 TDs (with 17 overlapping with Experiment 1) and 6 IDs participated in Experiment 2 (Actions experiment). However, one TD participant was not included in further analysis in Experiment 2 due to excessive head motion (see below)[13]. Resting-state imaging data were also collected for all participants. Five of the TDs were excluded from further analysis due to excessive head motion (see details below). Therefore, data from the remaining 17 TDs and all 9 IDs were included in the subsequent analysis.

None of the participants reported a history of psychiatric or neurological disorder. All experimental protocols and procedures were approved by the Institutional Review Board of Georgetown University (or Harvard University, for IDs 8 and 9) in accordance with the Declaration of Helsinki. Written informed consent was collected from each participant, and they were compensated for their time.

### Experimental Design and Procedures

#### Experiment 1: Motor Mapping Experiment

Experiment 1 (Motor Mapping Experiment, block design fMRI experiment) focused on motor responses of different body parts. In this experiment, all participants were asked to perform simple movements with different body parts, including the mouth, abdomen, left and right shoulders, left and right feet, and left and right hands (just for typically developed participants), similar to the previous study [5]. Each body part was moved (simple flexing/contraction movement) in separate blocks in randomized order according to an auditory cue (metronome).

Each run began with a baseline period, followed by alternating movement and rest blocks. fMRI data of two runs from participants ID1-ID7 were collected at Georgetown University using a block design with 6.256 s (8 TRs) movement and 6.256 s rest. Four flex and relax movements were performed in each block at a frequency of 0.48 Hz.

fMRI data of three runs from participants ID8 and ID9 were collected at Harvard University using a similar block design with 6 s (3 TRs) movement and 6 s rest [5]. Four flex and relax movements were performed in each block at a frequency of 0.66 Hz.

#### Experiment 2: Actions Experiment

Experiment 2 (Actions Experiment, slow event-related fMRI experiment) focused on effector-specific brain activity (hands and feet in TDs, feet in IDs) during action execution [13]. In the Actions experiment, all participants were asked to perform four kinds of actions (grasping, drawing, writing, and simple tool-use). All the actions are grouped together for our analyses here, to examine body-part responses for goal-directed actions. Participants performed the actions with their right hands and feet (only feet in IDs), except ID6, who completed the actions with her dominant left foot. Due to technical considerations, each run corresponded to action execution by only one effector (846TRs per run). The first four typically developed participants had an additional experimental condition, which was later omitted, resulting in runs of 950TRs. Each run began with a baseline period of 7.8 seconds (10TRs), followed by pseudorandomized grasping, drawing, writing, and tool-use trials (six grasping, eight drawing, four writing, and twelve tool-use trials per run). Each trial began with an auditory instruction of the action. Following the instructions, participants had 4.7 seconds (6TRs; action planning) before hearing the first sound cueing them to begin executing the action (action execution). Only this stage of action execution is analyzed for our purposes here. Participants then had 5.5 seconds (7TRs) to complete the action and were instructed to hold the action’s end position until they heard a second sound, cueing them to return to the resting position (return period). During the return period, participants would either return the object to its starting position on the board (hand trial) or drop it (foot trial), as most control participants could not quickly return the object with their feet (1.6 seconds, 2TRs). Trials were spaced by 8.6 seconds (11TRs). Separate practice sessions were carried out prior to the in-scanner experiment to familiarize the participants with the trial timing and object manipulation instructions.

For each grasp action, participants reached and grasped a spatula or a crayon without lifting the object. For each draw action, participants reached and grasped a crayon and drew a triangle or a diamond. For each writing action, participants reached and grasped a crayon and wrote the letters “n”, “g”, or “s”. For each simple tool-use action, participants reached and grasped a spatula and used it to spread balloons to the left or the right (mimicking the action of frosting a cake or spreading butter on toast). As our goal here was to compare the effector-specific activity in SII in TDs and IDs, we averaged brain activity across the four different actions.

#### Resting-state fMRI

Resting state fMRI data were also collected from all 22 TDs participants and 9 IDs participants. Resting-state data were collected while the participants lay supine in the scanner without any external stimulation or task, and with their eyes closed.

### Functional Imaging

For all TDs participants, and seven IDs participants (ID1-ID7), functional imaging data of Experiment 1 (Motor mapping Experiment), Experiment 2 (Actions Experiment), resting-state data and anatomical images were collected in a Siemens Prisma-Fit 3T scanner at the Center for Functional and Molecular Imaging at Georgetown University Medical Center using a 32-channel head coil. Functional task data were acquired with a T2-weighted segmented gradient echo-planar imaging sequence (repetition time [TR] = 782 ms, echo time [TE] = 35 ms, flip angle [FA] = 52°, FOV=208 × 200 mm, matrix = 104 × 100, slice thickness = 2 mm, distance factor = 0%, voxel size = 2 × 2 × 2 mm^3^). The whole brain was covered by 72 interleaved slices, transversally oriented. 6-minute (767 TRs) resting state fMRI data was collected using a multiband echo-planar imaging (EPI) sequence with the following parameters: TR = 782 ms, TE = 35 ms, flip angle = 52°, FOV = 208 x 200, matrix = 104 × 100, slice thickness = 2 mm, distance factor = 0%, voxel size = 2 × 2 × 2 mm^3^. The whole brain was covered by 72 interleaved slices, transversally oriented.

For two IDs participants (D8 and D9), functional imaging data of Experiment 1 (Motor Mapping Experiment), resting-state fMRI data and anatomical images were obtained in a Siemens Tim Trio 3T scanner at the Center for Brain Science at Harvard University. Experiment 1 was collected with a 6-channel birdcage head coil (for details, please see supplementary information in [5]. Resting state functional imaging data were obtained using a 32-channel head coil. A total of 400 volumes (6 minutes) of multiband echo-planar imaging (EPI) sequence was acquired with the following parameters: TR = 1500 ms, TE = 28 ms, flip angle = 75°, FOV = 216 × 216, matrix = 108 × 108, slice thickness = 2 mm, voxel size = 2 × 2 × 2 mm^3^. A total of 68 axial slices were acquired with a slice spacing of 2.2 mm, covering the whole brain.

### Anatomical imaging

For all TDs participant and seven IDs (ID1-ID7), a T1-weighted anatomical reference volume was collected using a 32-channel head coil with an MPRAGE sequence (TR = 1900 ms, inversion time [TI] = 900 ms, FA = 9 degrees, FOV = 256 × 256mm, distance factor = 50%, 1 mm isotropic resolution). The whole brain was covered in 176 ascending slices in a sagittal orientation.

For two IDs participants (ID8-ID9), a T1-weighted anatomical reference volume was collected using a 32-channel head coil with a MPRAGE T1-weighted sequence with following parameter: TR = 2530 ms, TE = 1.64 ms, TI = 1200 ms, flip angle = 7°, FOV = 256 × 256 mm, 1 mm isotropic resolution.

### Data Analysis

AFNI [72], SUMA [73], FreeSurfer [74], Matlab (R2023a, MathWorks), R (4.5.0)[75], and JASP (Version 0.19.3.0) were used for data analysis.

#### Task-based fMRI data preprocessing

For the functional imaging data, preprocessing was carried out in AFNI [72]. The preprocessing procedure included the following steps: removal of spikes, slice-timing correction, head-motion correction by realignment of functional volumes, alignment of functional images to the structural image; non-linear transformation to Montreal Neurological Institute (MNI) 152 template, spatial smoothing with 4-mm full-width at half maximum (FWHM) Gaussian filter; and voxel-wise normalization of time series.

#### General linear modeling (GLM)

For whole-brain analysis in task-based Experiments 1 and 2, we performed voxel-wise mixed-effects general linear model analysis in AFNI [72]. In Experiment 1 (Motor Mapping experiment), left and right hands (for TDs), feet, shoulder, mouth, and abdomen were included as regressors in the analysis. In Experiment 2 (Actions experiment), grasping, drawing, writing and tool-use action execution stages with the right hand (for TDs) and right foot were included as regressors in the analysis. Beta values were estimated using a GLM procedure, convolved with the standard Hemodynamic Response Function (HRF), and fit using a voxelwise generalized least squares (GLS) with an autoregressive moving average-like noise model [72]. Due to our focus on the compensatory use of the feet, and as most IDs were dominantly right-footed, we used the movements of the right hand (in TDs) and foot for further examination. Simultaneously, covariates of no interest (e.g., six rigid-body motion parameters and their temporal derivatives, signals of ventricular cerebrospinal fluid (CSF) tissues defined using FreeSurfer [74] were also included in GLM analysis, along with a local white-matter nuisance regression. In Experiment 1 (Motor Mapping Experiment), volumes with framewise motion (Euclidean norm of motion derivatives) larger than 0.5 mm or with an outlier fraction larger than 0.05 were censored from the analysis. In Experiment 2 (Actions experiment), where more body movement was involved due to performing more complicated actions in the scanner than Experiment 1, volumes with framewise motion (Euclidean norm of motion derivatives) larger than 1 mm or with an outlier fraction larger than 0.05 were censored from the analysis. One TD participant in Experiment 2 was excluded for further analysis due to excessive motion with more than 50 TRs (corresponding to approximately 5-6% TRs) censored in each run. In Experiment 1 (Motor Mapping Experiment), we localize the hand area by contrasting responses for the right hand and the other body parts (right foot, right shoulder, abdomen, and mouth) for TDs. Activation maps were corrected for multiple comparisons at p < 0.05 using the cluster-level correction based on Monte Carlo stimulations implemented in AFNI [72, 76]. To discover the brain reorganization in IDs, we inspected the body part selectivity in the hand-selective region (as defined in TDs) by contrasting each body part activity and the other three body part activities. Given the limited sample size in the IDs group (N = 9), due to the rarity of congenital bilateral hand absence, group-level body selectivity analysis was performed using a mixed-effects linear mixed model in AFNI, including the scanning location as a covariate to account for potential scanner and sequence effects. For winner-takes-all analyses (Fig. S1), each voxel was then labeled with the condition that produced the highest average activation. This analysis was limited to the SII area, defined by architecture area of parietal operculum1 (OP1) [27].

In experiment 2 (Actions Experiment), hand and foot activity in SII were extracted for goal-directed action execution in TDs, and foot activity was extracted in IDs. Given the limited sample size in the IDs group (N = 6), we applied a bootstrapping test to assess differences between hand or feet activity in TDs and feet activity in IDs using R [75]. This is a non-parametric resampling approach that estimates the sampling distribution of a statistic by repeatedly resampling (with replacement) from the observed data. Significance was determined by nonparametric bootstrapping (10,000 iterations). The difference was considered significant when the 95% bootstrap confidence interval (CI) did not include zero.

#### Resting-state fMRI data analysis

For resting-state data, preprocessing was carried out using AFNI [72]. Preprocessing steps included removal of spikes, slice-timing correction, head-motion correction by realignment of functional volumes, alignment of functional images to the structural image; non-linear transformation to Montreal Neurological Institute (MNI) 152 template, spatial smoothing with 4-mm full-width at half maximum (FWHM) Gaussian filter; and voxel-wise normalization of time series. Nuisance regression and temporal filtering included six rigid-body motion parameters and their derivatives, local white-matter signal, and the first three principal components from the lateral ventricular CSF ROI defined using FreeSurfer [74]. Given resting-state functional connectivity analyses’ sensitivity to motion, volumes with framewise motion larger than 0.2 mm or outlier fraction larger than 0.05 were censored.

Residual signals after nuisance regression were used for seed-based functional connectivity analysis. The hand-or foot-selective left primary sensorimotor cortex (SM1) and hand-selective left secondary somatosensory cortex (SII), defined in the Motor Mapping Experiment in TDs, were seeds in this functional connectivity analysis.

Repeated-measure Bayesian ANOVA with group (TDs, IDs) as group factors, and connectivity (connectivity between handSM1 and SII, connectivity between foot SM1 and SII) were applied to test how congenital hand absence shapes functional connectivity among the somatosensory cortices, with the scanning location as a covariate to account for potential scanner and sequence effects. This was calculated in JASP (Version 0.19.3.0).

## Competing financial interests

The authors declare no competing financial interests.

## Author Contributions

Liu. Y., and Striem-Amit. E., designed the study. Martinez-Addiego. F., and Liu. Y., performed the research. Deng. Z., analyzed the data. Striem-Amit. E., supervised the study. Deng. Z., wrote the draft of the manuscript. Striem-Amit. E., Martinez-Addiego. F., and Liu. Y. edited the manuscript. All authors have read and approved the final version of the manuscript, and agree with the order of presentation of the authors.

## Acknowledgments

We thank the participants who took part in our experiment. We are grateful to Gilles Vannuscorps and Alfonso Caramazza for providing and contributing to the Harvard dataset that supported part of the analyses reported here. This work was supported by the Edwin H. Richard and Elisabeth Richard von Matsch Distinguished Professorship in Neurological Diseases (to E.S.-A.) and a Shanghai Youth Science and Technology Innovation Plan (22YF1454200 to Y.L.).

## Data and materials availability

Data that support the main findings of this study, as well as the source maps for the figures data, have been deposited in OSF https://osf.io/h7zjv/overview.

The data and materials will be deposited into a public database, and there will not be any restrictions on data availability.

## Notes

### Competing Interest Statement

The authors have declared no competing interest.

