## Supplemental Figure 1 for "From Hands to Feet: Experience-Driven Plasticity in Secondary Somatosensory Cortex in People Born without Hands"

Supplementary information

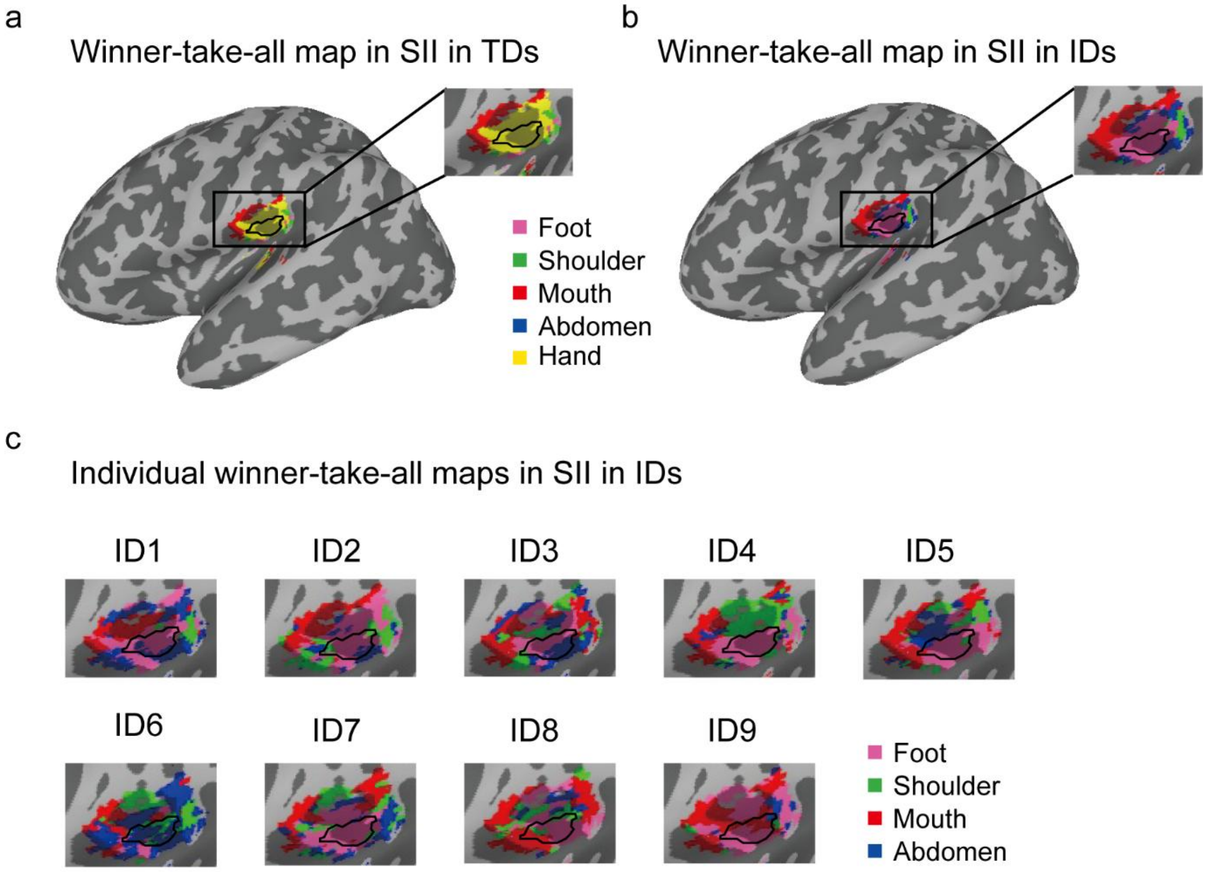


**Fig. S1: Mapping reorganization in individuals with upper-limb dysplasia (a)-(b)** winner-take-all map of typically developed individuals (TDs) and people born without hands (IDs) in the secondary somatosensory cortex (SII) confirms the takeover of the hand-selective area (From **Fig. 1a**; outlined in black) by the foot in IDs. The analysis was confined to SII, defined as the cytoarchitectural area parietal operculum1 (OP1) [1]. **(c)** individual winner-take-all maps of people born without hands (IDs) in the same SII region (OP1). Despite individual differences, in all but one ID participant (ID6), the area caudal and inferior to the mouth preference, where the hand preference would be, is mainly taken over by the foot [2, 3].
